# Molecular determinants of SEC61A1 mutant R236C in two patient-derived cellular platforms

**DOI:** 10.1101/2023.12.05.570221

**Authors:** Matthias Weiand, Vanessa Sandfort, Oksana Nadzemova, Robert Schierwagen, Jonel Trebicka, Iyad Kabar, Hartmut Schmidt, Andree Zibert

## Abstract

**Background & objectives:** SEC61A1 encodes a central protein of the mammalian translocon machinery residing in the ER. Mutations of *SEC61A1* are implicated to result in various severe diseases. Recently, mutation R236C was identified in patients having autosomal dominant polycystic liver disease (ADPLD), which originates from defects of cholangiocytes in the liver. The molecular phenotype of R236C was assessed.

**Methods:** Two cellular platforms were established from the same cell source of an ADPLD patient by different methodology. Cells were immortalized by retroviral transduction of an oncogene (UCi) or reprogrammed to induced pluripotent stem cells (iPSC) that were differentiated to cholangiocyte progenitor-like cells (CPLC). UCi and CPLC were subjected to several analyses of molecular pathways that were previously associated with development of polycystic disease.

**Results:** UCi displayed markers of epithelial cells, while CPLCs expressed typical markers of both cholangiocytes and hepatocytes. Cells encoding R236C showed a stable, continuous proliferation in both platforms, however growth rates were reduced as compared to wildtype control. Autophagy, cAMP synthesis, and secretion of important marker proteins were reduced in R236C-expressing cells. In addition, R236C induced increased calcium leakiness from the ER to the cytoplasm. Upon oxidative stress, R236C led to a high induction of apoptosis and necrosis.

**Conclusion:** Although the grade of aberrant cellular functions differed between the two platforms, overall the molecular phenotype of R236C was shared suggesting that the mutation, regardless of the cell type, has a dominant impact on disease-associated pathways that cause ADPLD. Our approach may be valuable to decipher the molecular pathways of other rare genetic diseases using patient-specific cell models.

## Introduction

Polycystic liver disease (PLD) belongs to a group of cystic disorders, characterized by fluid-filled hepatic cysts, typically caused by the mutation of a distinct genetic loci and affects the function and integrity of the liver [1]. PLD is associated with formation of cholangiocyte-derived hepatic cysts that progressively replace liver tissue [2]. Polycystic liver cysts can coexist with cysts of the kidney in autosomal dominant polycystic kidney disease (ADPKD) (prevalence ~1:400) and autosomal recessive polycystic kidney disease (ARPKD) (prevalence ~1:20,000). In contrast, only few, clinically not relevant kidney cysts are usually observed in autosomal dominant polycystic liver disease (ADPLD) which is rare (prevalence ~1:100,000) [3]. ADPLD can be associated with severe disease of mostly adult onset. As no approved drug is efficient, PLD can only be cured by liver transplantation.

Mutations of polycystin-1 and polycystin-2 (*PKD1* and *PKD2*, respectively) account for most ADPKD, whereas mutations of *PKHD1* and less frequently *DZIP1L* account for most ARPKD [4]. Seven mutations, predominantly affecting the processing and quality control of glycoproteins in the endoplasmatic reticulum (ER) have been identified to cause ADPLD (*PRKCSH, SEC63, ALG8, SEC61B, GANAB, LRP5*, and *PKHD1*) [3]. However, for a large portion of PLD patients the disease-causing mutation is unknown [5]. Recently, a novel mutation R236C encoded by *SEC61A1* was identified in two patients having ADPLD [6].

The trimeric SEC61 mammalian complex consists of the proteins SEC61A1, SEC61B and SEC61G. The protein SEC61A1 comprises ten transmembrane helices which and several cytosolic and lumenal loops that are proposed to form the central channel [7]. The complex functions as a central part of the translocon machinery facilitating the co- or posttranslational signal peptide-dependent polypeptide transport into the lumen of the ER for endo- and exosomal transport of many proteins [8, 9]. The SEC61 complex is highly regulated by allosteric effectors [10]. Recently, the pore of the SEC61 complex was shown to mediate the passage of small molecules and ions, importantly calcium (Ca) [11]. The open conformation of the SEC61 complex allows the passive calcium efflux from the ER to the cytoplasm. The complex is also highly related to processes of the unfolded protein response (UPR) as well as autophagy and apoptosis [8, 12, 13].

Serving as the entry point of newly synthesized peptides and strongly connected to various cellular key functions, SEC61 is highly conserved and expressed in all nucleated cells [14]. Knockdown or mutation were shown to cease cell proliferation demonstrating the essential role of SEC61 for cellular integrity [15]. Consequently, sub-lethal malfunction of SEC61 results in a variety of different disease, which were proposed to form an own entity and termed SEC61 channelopathies [10].

To date, only a few mutations of *SEC61A1* have been identified in patients, importantly V67G, V85D, Q92R, T185A, and E381*. However, none of the mutations was associated with PLD. As mutation-specific animal models are missing, functional analysis of a given *SEC61A1* mutation is frequently confined to the study of commercial cell lines [15–17]. However, such expression systems lack the genuine genetic background of the patient and other organ-specific constraints. In addition, primary cells encoding mutations that cause channelopathies, are difficult to obtain from rare patients. Such procedures frequently afford invasive methodology for isolation of cells and primary cells usually do not proliferate long-term.

We have previously reported that primary cells derived from urine (UCs), representing a non-invasive cell source, can be immortalized via retroviral expression or reprogrammed to induced pluripotent stem cells (iPSCs) followed by liver-specific differentiation [18]. In the present study, basic molecular functions of SEC61A1 mutation R236C were studied using this methodology of different cellular platforms. UCs were either immortalized (UCi) or differentiated to cholangiocyte progenitor-like cells (CPLC) from iPSCs. The molecular phenotype of the SEC61A1 mutation was assessed for several biochemical hallmarks of polycystic diseases, like autophagy, apoptosis, intracellular Ca-homeostasis, and ER-related protein biosynthesis.

## Material and Methods

### Ethics statement

Urine samples were obtained in accordance with the ethics committee of the Universitätsklinikum Münster. Participants were informed regarding the use of data and written informed consent was obtained from all individuals enclosed in the study.

### Isolation of urine-derived cells

Cells were harvested from urine samples as reported before (recruitment period: 08/10/2020– 11/09/2020) [19]. Cultivation of cells was in DMEM/F-12 (Gibco), 10 % FCS (Gibco), 1 % PenStrep (Gibco), 1x MEM non-essential amino acid solution (NEAA, Sigma), 1x GlutaMAX-I (Gibco), 0.1 mM 2-mercaptoethanol (Gibco) and SingleQuot Kit CC-4127 REGM (Lonza). Subconfluent primary cells were expanded for and frozen for experiments.

### Retroviral transduction

Human embryonic kidney HEK293T cells were maintained in DMEM/F12 (Gibco) with 10 % FCS (Gibco) and 1 % PenStrep. For generation of retroviral supernatants, HEK293T cells were transfected with plasmid pLXSN16E6E7 encoding the human papilloma virus HPVE6E7 gene (Addgene #52394) as described [18]. Urine cells following retroviral transduction were cultivated in UC medium (DMEM/F12, 10 % FCS, 1 % L-Glu, 1 % PenStrep).

### Induced pluripotent stem cells

Establishment of iPSCs was performed as reported before [18]. 0.5 x 10^6^ primary urine cells were reprogrammed using the Amaxa Basic Nucleofector Kit (Lonza) according to the protocol of the manufacturer. 1 μg of each plasmid pCXLE-hOCT3/4-shp53-F (Addgene #27077), pCXLE-hSK (Addgene #27078), and pCXLE-hUL (Addgene #27080) was used. After nucleofection, the cells were transferred to a Matrigel (Corning) coated 6 well. After 24 hours, the UC medium was replaced by mTeSR-1 (Stemcell Technologies), supplemented with 1% PenStrep, and changed every second day. After about two weeks the emerged iPSC colonies were picked and transferred to a new Matrigel-precoated 6 well plate. The mTeSR-1 cell medium (was replaced on a daily basis thereafter and the iPSC colonies were subcultured every seven days using 1 U/ml Dispase (Stemcell Technologies). During cultivation, colonies with atypical morphology were removed by scraping.

### Differentiation of cholangiocyte progenitor-like cells

For cholangiocyte progenitor-like cell differentiation a protocol used before for hepatocyte-like cells was slightly modified [18]. Briefly, iPSCs clusters instead of single cells were seeded to Matrigel-precoated wells in order to achieve an almost confluent cell layer at the next day. The medium was changed daily for the next three days to DMEM/F12 supplemented with 100 ng/ml recombinant Activin-A (R&D Systems), 100 ng/ml fibroblast growth factor-2 (FGF2, Peprotech), 50 ng/ml recombinant human Wnt3a (R&D Systems; day 1 and 2), 1 mM L-Glutamine (L-Glu, Sigma). KnockOut SR Xenofree CTS (KSR, Gibco) was added at 0.2 % (day 2) and 2 % (day 3). For the next 8 days (days 4-11), cells were cultivated in DMEM/F12 supplemented with 10 % KSR, 1 mM NEAA, 1 mM L-Glu, 1 % dimethyl sulfoxide (DMSO, Sigma), and 100 ng/ml hepatocyte growth factor (HGF, Peprotech). The maturation step, was prolonged up to day 15. 0.1 μM dexamethasone medium was added on day 12 to the medium (DMEM/F12, 10 % KSR, 1 mM NEAA, 1 mM L-Glu, 1 % DMSO) without any further medium changes at day 14 and day 15.

### PCR

Total RNA isolation was performed by RNeasy Mini Kit (Qiagen) and first strand synthesis was accomplished using SuperScript™ III First-Strand Synthesis SuperMix (Thermo). For real time PCR, Takyon ROX SYBR Master Mix blue (Eurogentec) was used with a 0.4 µM final primer concentration (Supplementary Table S1). qPCR was conducted on QuantStudio 7 Pro device (Thermo). Ct values were normalized to GAPDH and evaluated using the comparative Ct method (2^-ΔCt^).

### Flow cytometry analysis

For antibody staining, cells were fixed with 2 % Histofix (Roth) for 30 min and permeabilized by 0.05 % Triton X-100 (Thermo) for 15 min. After blocking in 3 % BSA solution (Sigma) for a minimum of one hour, cells were incubated with antibody in a 1 % BSA solution (Supplementary Table S2). For autophagy, trypsinized cells were stained with CYTO-ID according to the protocol of the CYTO-ID® Autophagy detection kit (Enzo) using 5 % FCS. CYTO-ID staining was for 45 min. For apoptosis, 5 mM or 10 mM H_2_O_2_ (AppliChem) was incubated with the cells (24 well) in cell culture medium. Incubation time was as described in the figure legend. Cells were then trypsinized and centrifuged (400 g, 5 min) including the respective cell culture supernatants. The cell pellet was resuspended in 50 µl of the Annexin V Apoptosis Detection Kit/7-AAD solution (BioLegend) and incubated for 15 min. Flow cytometry analysis was accomplished in a CytoFlex Flow Cytometer (Beckman Coulter).

### Immunocytochemistry

For immunocytochemistry, the cells were fixed using 4% Histofix (Roth) for 30 min and permeabilized by 0.05 % Triton X-100 for 15 min. Blocking (3 % BSA in PBS) and antibody staining (1 % BSA in PBS) was conducted overnight (Supplementary Table S2). Following DAPI (Sigma) counterstaining, photographs were taken using an Olympus CKX41-X10 microscope with CellSens Standard 1.61 (Olympus) imaging software.

### Western blot

Lysates were prepared using RIPA buffer (60 mM tris-HCl, 150 mM NaCl, 2 % Na-deoxycholate, 2 % Triton X-100, 0.2 % SDS and 15 mM EDTA) in the presence of protease inhibitors (Roche, Complete Mini, EDTA-free). Blotting was conducted using the Trans-Blot Turbo Transfer System (Biorad). The PVDF membranes (Biorad) were blocked for at least one hour in TBST-T (20 mM Tris, 150 mM NaCl, 0.05 % Tween-20) with 5 % skimmed milk (Roth). For primary antibody staining, membranes were incubated in TBST-T with 0.5 % skimmed milk overnight. After secondary antibody staining (1 h), membranes were washed and Clarity Western Blot ECL Substrate (Biorad) was added. Imaging was accomplished by the Fusion Solo S imaging system (Vilber). ImageJ 1.53k analysis software was used for densitometric quantification using GAPDH as reference.

### Intracellular calcium measurement

For calcium measurement, cells grown on a 24 well plate were washed and incubated in 200 µl of HBSS buffer (10 % FCS, 0.04 % F-127, 2 mM Probenecid, 1.5 mM CaCl_2_, 10 mM HEPES). Two µl Fluo-4 (AM, Invitrogen; 0.45 mM in DMSO) were added. Incubation was for 45 min. After this, medium was changed to DMEM/F12 (5 % FCS, 0.04 % F-127, 2 mM Probenecid). 1 mM Thapsigargin (Hello Bio) was added to a final concentration of 1 mM. A video was recorded with the Olympus software before and after addition of TG. The recorded videos were analyzed with ImageJ 1.53k as follows. The video was converted to greyscale. 50 cells were selected randomly and marked as regions of interest. 10 cell free sections served as background. The fluorescence intensity for every region of interest was determined. From these values the mean fluorescence intensity of the background was subtracted. Values were normalized to the mean fluorescence before TG addition (baseline).

### Protein expression

Human PreAlbumin (TTR) ELISA Kit (abcam) and cAMP Parameter Assay Kit (R&D) were performed as recommended by the supplier. The Human VEGF Magnetic Luminex® Performance Assay (R&D) was used for determination of VEGF. Total protein determinations were accomplished by Bradford assay (Biorad) and used for normalization. Photometrimetic measurements were done by a Tecan SPARK microplate reader or the Luminex 200 system (R&D).

### Statistical analyses

Statistical analyses were performed by Mann-Whitney U, Student’s t-test and Wilcoxson matched-pairs signed rank test using GraphPad Prism (v6.01, GraphPad Software) software. Data are given as mean ± standard error of mean (SE).

## Results

### Establishment of SEC61A1 R236C in two cellular platforms obtained from ADPLD patient

Two patient-specific cellular platforms, representing urinary epithelial cells and iPSC-derived cells [18], were used throughout the study to model the molecular consequences of a R236C [6]. Urine cells (UCs) derived from an ADPLD patient encoding R236C were subjected to retroviral gene transfer for immortalization (UCi) or reprogrammed to iPSCs followed by differentiation to cholangiocyte progenitor-like cells (CPLC) (Figure 1A). UCs from an age-matched, disease-free individual expressing SEC61A1 wild type (WT) were subjected to identical methodology and used as a control. UCi_R236C_ and UCi_WT_ showed continuous proliferation for more than 12 months (data not shown) and presented a typical smooth-edged shape and cobblestone-like morphology (Figure 1B). To assess any differences between UCi_R236C_ and UCi_WT_, the expression of epithelial (*CK7* and *CLDN1*), renal (*L1CAM* and *SNAI2*), and fibroblast marker (*FN1*), that are commonly found [19], was performed by RT-qPCR analysis (Figure 1C). Flow cytometry analysis revealed high (>95.0 %) expression of epithelial markers CD13, CD29, and CD166 in UCi_R236C_ and UCi_WT_, whereas expression of mesenchymal stem cell markers CD71 and CD105 were increased (>20.0 %) in UCi_R236C_ as compared to UCi_WT_ (Figure 1D).

**Figure 1:**
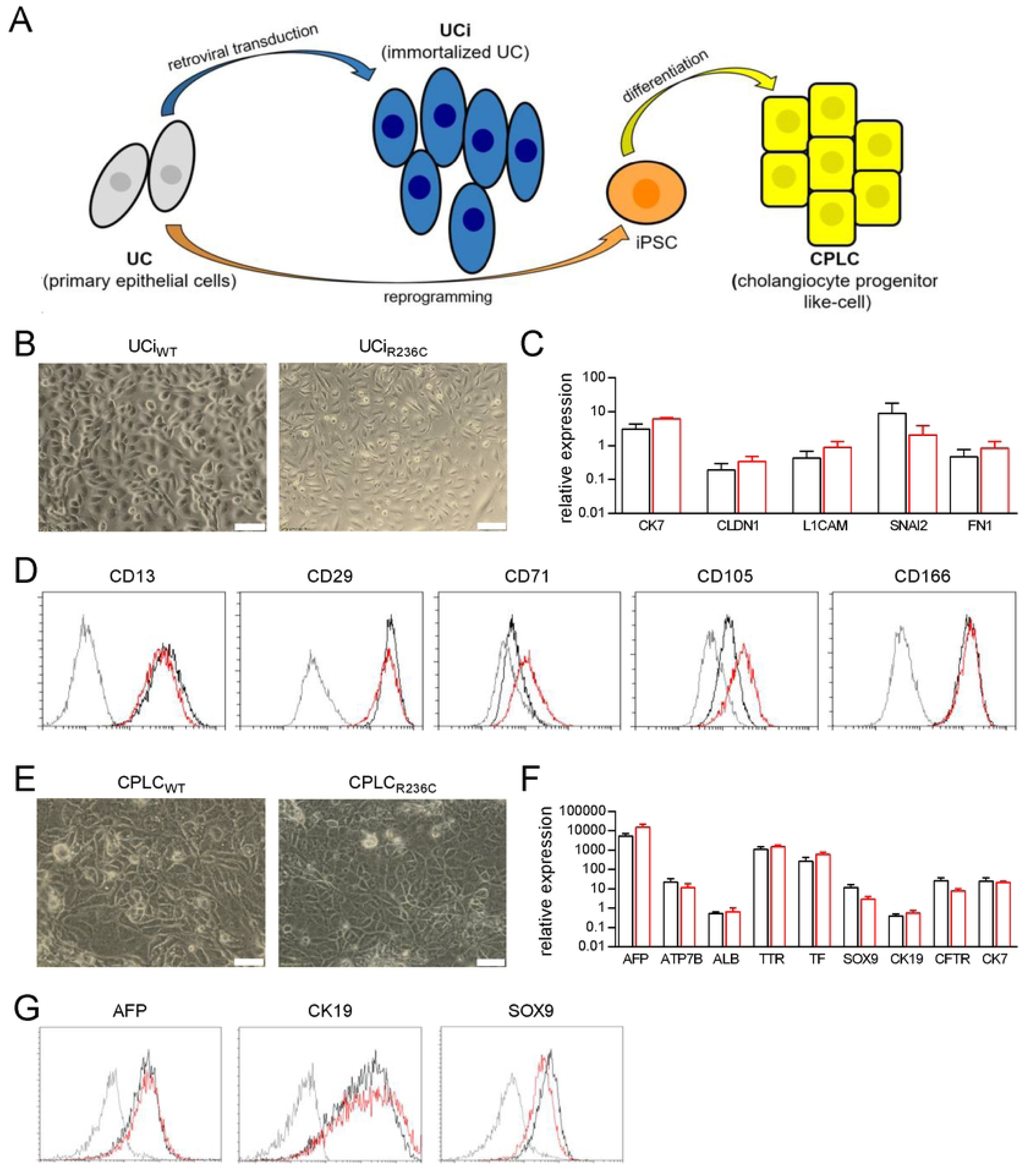
SEC61A1 mutant R236C can be established in UCi and CPLC platforms. (A) Schematic overview of UCi and CPLC generation. (B) Typical bright field image of UCi derived from wildtype (WT) and R236C. Scale bars, 100 μm. (C) Marker gene expression of immortalized UCi as assessed by RT-qPCR analysis. Data were normalized to GAPDH of primary UCs. Mean and SE are shown (n=6). *P < 0.05. (D) Flow cytometry analysis of UCi. Isocontrol (shaded line), UCi_R236C_ (red) and UCi_wt_ (black) are shown. One typical experiment of three is shown. (E) Bright field image of CPLCs. One typical experiment is shown. Scale bars, 100 μm. (F) Marker gene expression of CPLCs as assessed by RT-qPCR analysis. Data were normalized to GAPDH of biliary reference cell line HepaRG. Mean and SE are shown (n=8). *P < 0.05. (G) Flow cytometry analysis of CPLCs. Isocontrol (shaded line), CPLC_R236C_ (red) and CPLC_wt_ (black) are shown. One typical experiment of three is shown.

UCs were also reprogrammed to iPSCs and differentiated to CPLCs. iPSCs showed similar cell morphology and marker expression of typical stem cell markers (NANOG, OCT4, SOX2, and SSEA-4) suggesting that R236C does not impose reprogramming (Supplementary Figure S1). After differentiation to CPLCs, cells exhibited a rhomboid shape indicative of a cholangiocyte-like differentiation [20] (Figure 1E). Immunohistochemical analysis revealed the expression of hepatocyte markers AFP, ALB, and HNF4α (Supplementary Figure S2). In RT-qPCR analysis (Figure 1F), CPLC_R236C_ and CPLC_wt_ showed similar levels of hepatocyte markers (*AFP, ATP7B*, *ALB, TTR,* and *TF)* and cholangiocyte markers (*SOX9, CK19, CFTR,* and *CK7).* Expression of *AFP, ATP7B*, *TTR, TF*, *SOX9, CFTR* and *CK7* was high, whereas *ALB* and *CK19* expression was in the same range as in the reference cell line. Immunostaining of CPLC_R236C_ and CPLC_wt_ also suggest the presence of hepatocyte and cholangiocyte markers (Figure 1G). Overall, the coexpression of cholangiocyte and hepatocyte markers indicates that our reprogramming/differentiation protocol leads to a progenitor cell type resembling aspects of physiological liver organogenesis [21]. The genotype of R236C was maintained in tissue culture following the events of retroviral immortalization and iPSC establishment (Supplementary Figure S3). Of note, although immortalized and reprogrammed cells showed stable continuous proliferation, growth rates of UCi_R236C_ and CPLC_R236C_ were lower as compared to control suggesting that R236C might interfere with the progress of the cell cycle (Supplementary Figure S4).

### SEC61A1 R236C results in reduced autophagy and cAMP levels

SEC61A1 was previously reported to affect the level of autophagy [22]. Autophagy of R236C-expressing cells was investigated by Western blot using the analysis of LC3-II [23]. Level of LC3-II was reduced in UCi_R236C_ and CPLC_R236C_ relative to control (Figure 2A and Figure 2B, respectively) suggesting that autophagy is impeded by expression of R236C. In order to reassess autophagy by a different methodology, cells were also stained with CytoID, which specifically binds to autophagic compartments [24]. Flow cytometry analysis revealed a reduced mean fluorescence (MFI) in UCi_R236C_ and CPLC_R236C_ relative to control (Figure 2C and Figure 2D, respectively) corroborating that R236C impairs formation of autophagosomes.

**Figure 2:**
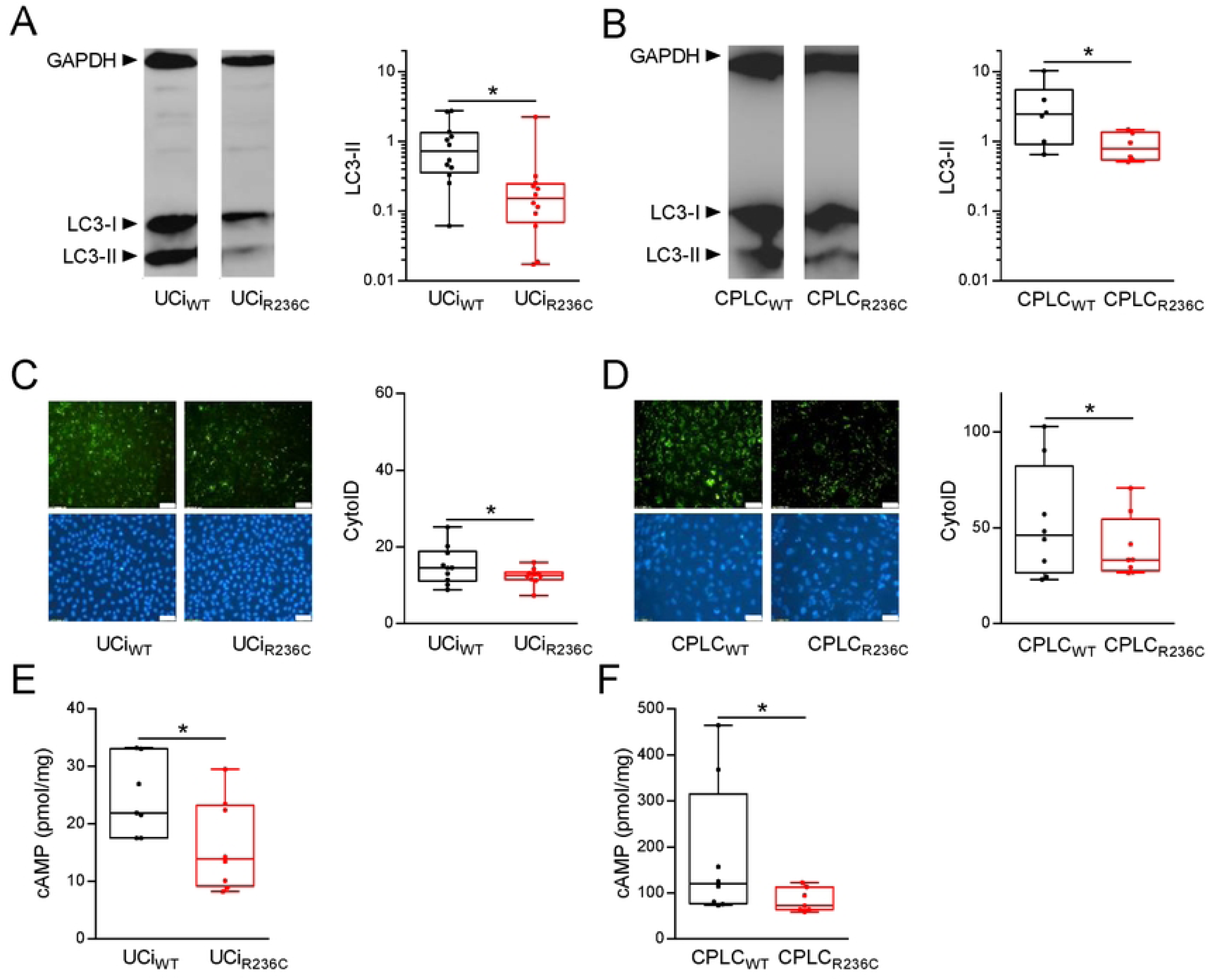
R236C results in reduced autophagy and decreased intracellular cAMP. (A) UCIs and CPLCs (B) were subjected to Western blot analysis using autophagy markers LC3-I and LC3-II. One typical example is shown. Box plots depict relative expression of LC3-II in UCis (n=12) and CLPCs (n=6) following densitometry analyses. GAPDH served for normalization. *P < 0.05. (C) UCis and CPLCs (D) were subjected to staining by CytoID. One typical fluorescence image is shown. Scale bars, 100 μm. Box plots depict MFI obtained after CytoID staining and flow cytometry analysis of UCi (n=10) and CPLC (n=8). *P < 0.05. (E) UCis and CPLCs (F) were subjected to intracellular cAMP measurements using cell lysates (n=7). *P < 0.05.

As cyclic adenosine monophosphate (cAMP) was reported to be associated to autophagy [25], the level of intracellular cAMP was determined. The cAMP level in UCi_R236C_ (Figure 2E) and CPLC_R236C_ (Figure 2F) was decreased relative to control indicating that cAMP-mediated intracellular signaling is downregulated in cells encoding R236C.

### SEC61A1 R236C stimulates induction of apoptosis

As autophagy and apoptosis are highly related [26], apoptosis was also studied. The question was addressed whether R236C modulates induction of apoptosis. In order to assess apoptosis, the response to oxidative stress was determined following exposition of cells to reactive oxygen species (ROS) and flow cytometry analysis of Annexin V (AV) and 7-AAD (AD). Different conditions of ROS exposure were used, since it was shown that UCs were more sensitive to ROS as compared to CPLCs (Supplementary Figure S5). A high induction of apoptotic cells (AV+) was induced in R236C-expressing cells upon exposure to ROS (Figure 3). Necrosis (AD+) was higher in UCi_R236C_ as compared to CPLC_R236C_ suggesting that R236C leads to induction of predominantly late apoptosis (AV+/7AD+) in urinary epithelial cells, whereas early apoptosis (AV+/7AD-) is predominantly induced in cholangiocyte-percursor cells. Of note, CPLC_R236C_ that were not challenged by ROS demonstrated a high level of apoptotic cells (AV+) as compared to UCi_R236C_, however such numbers did not reach significance as compared to control.

**Figure 3:**
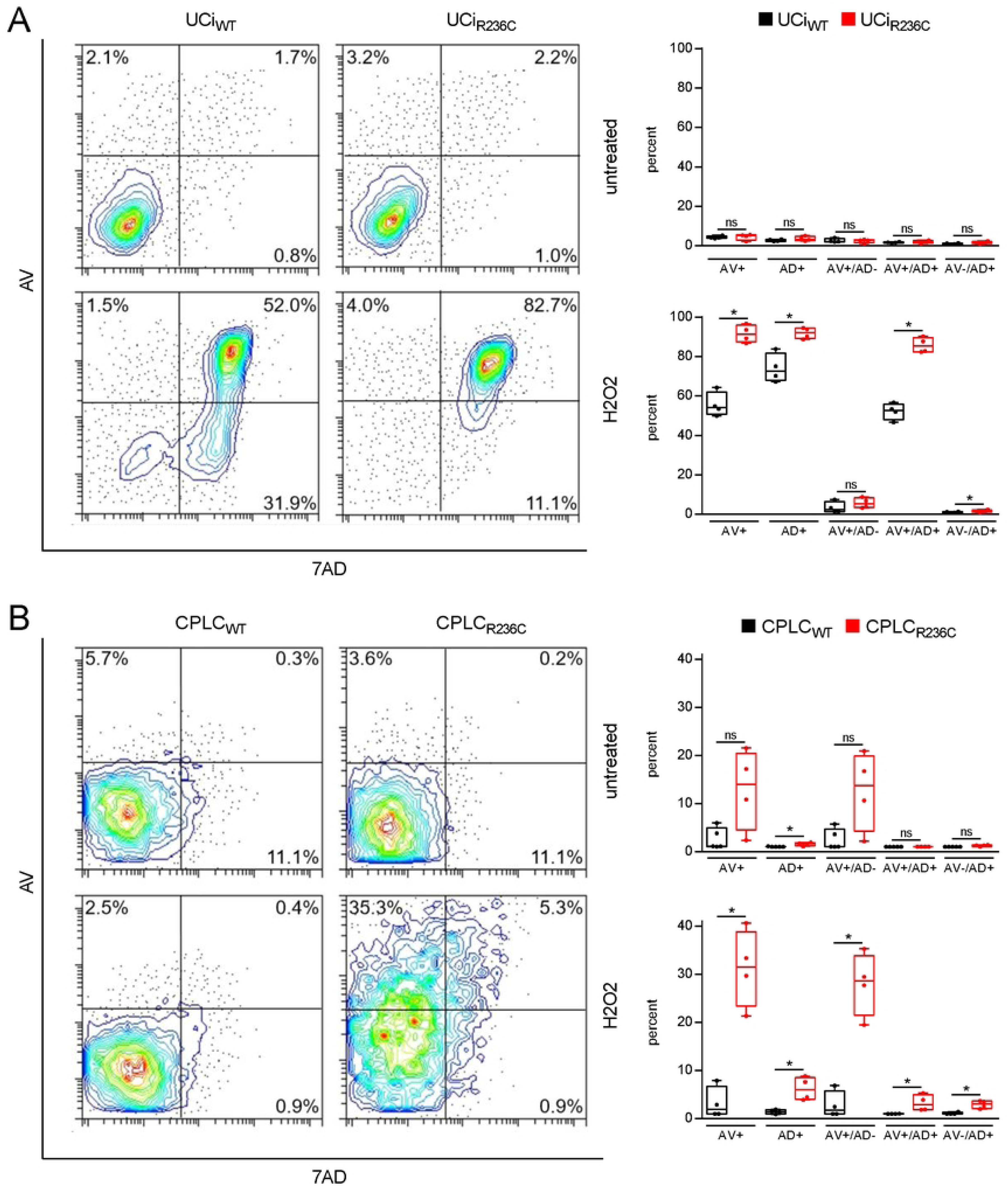
R236C increases sensibility to oxidative stress and induction of apoptosis. (A) Flow cytometry analysis of UCis and CLPCs (B) following staining by Annexin V (AV) and 7AAD (AD). Untreated control and cells after H_2_O_2_ exposure are shown. Treatment was extended from 3 h at 10 mM in UCi to 21 h at 5 mM in CLPC. Box plots depict percentages of cell populations obtained with UCi (n=10) and CPLC (n=8). *P < 0.05.

### SEC61A1 R236C results in increased calcium leakiness

The SEC61 complex was described as passive calcium channel that contributes to calcium release from the ER to the cytoplasm [27]. The question was addressed whether R236C affects intracellular calcium homeostasis. Thapsigargin (TG) treatment of Fluo-4 stained cells revealed that fluorescence intensity was reduced in UCi_R236C_ and CPLC_R236C_ relative to control (Figure 4A). Time-course analyses indicated that mean fluorescence intensity was reduced starting at ~40 seconds in UCi_R236C_ and CPLC_R236C_ relative to control (Figure 4B). In addition, the maximal fluorescence intensity at turning point (70s) was reduced in UCi_R236C_ and CPLC_R236C_ relative to control suggesting that R236C leads to increased calcium leakiness in both cellular platforms (Figure 4C).

**Figure 4:**
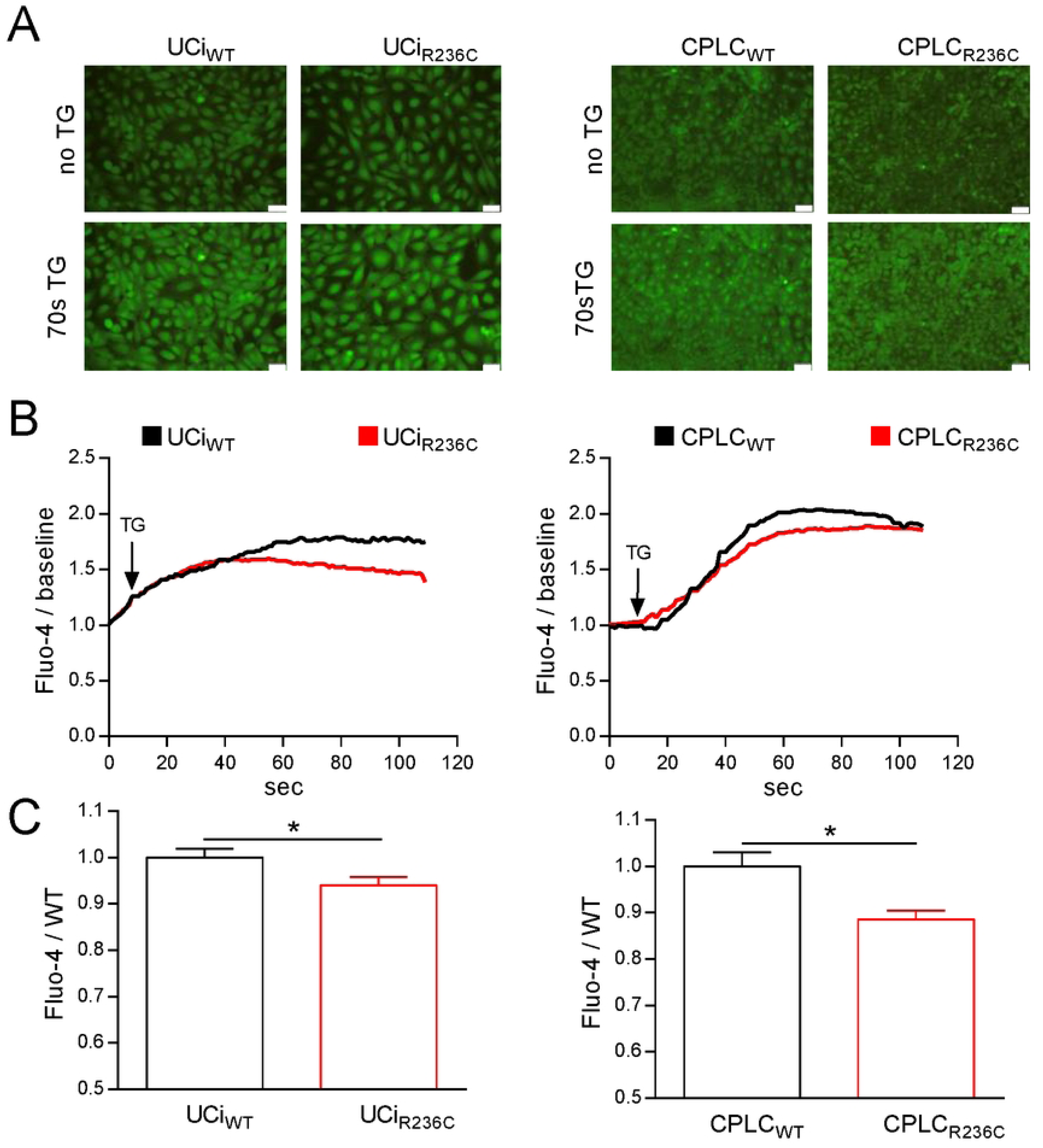
R236C causes calcium leakiness. (A) Fluorescence imaging obtained after incubation with Fluo-4. One typical image prior addition of thapsigargin (no TG) and after 70 seconds of TG are shown. Scale bars, 100 μm. (B) Mean calcium level over time is shown (n = 7). (C) Bar diagram indicates relative Fluo-4 fluorescence obtained after the curve reached the turning point (70 seconds). Mean and SE are shown (UCi: n=7). *P < 0.05.

### Sec61A1 R236C leads to lower protein secretion

The heterotrimeric SEC61 complex has a key role in translocation across the ER membrane for many proteins of the secretory pathway [8]. To assess the impact of R236C for protein translocation, the concentration of two proteins, VEGF and TTR, known to be dependent on SEC61 [28, 29] was determined in cell culture supernatants. The level of VEGF was decreased in UCi_R236C_ and CPLC_R236C_ relative to controls (Figure 5A). TTR, known as a typical hepatocyte marker [19], displayed a reduced concentration in CPLC_R236C_ relative to control (Figure 5B). In contrast, UCi did not secrete detectable levels of TTR corroborating the distinctness of the two cellular platforms.

**Figure 5:**
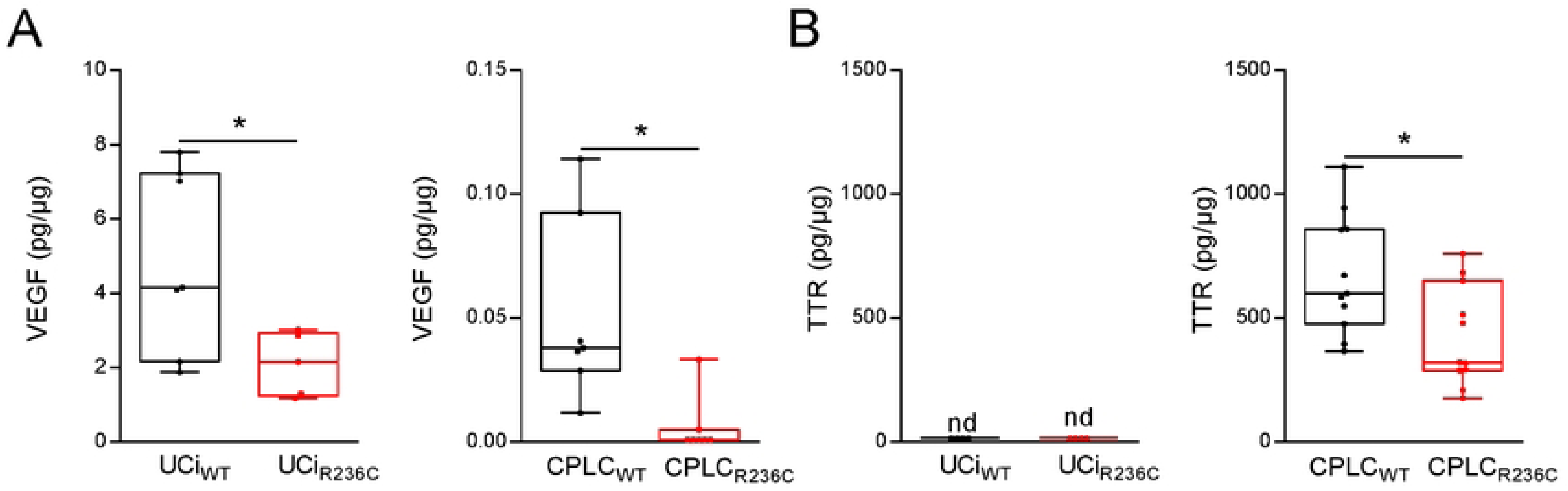
R236C impairs level of secreted proteins. (A) Secretion of VEGF and TTR (B) was determined in the cell culture supernatant of UCi (n=7) and CPLC (n=11). Total protein amount was used for normalization. *P < 0.05. nd, not detected.

## Discussion

We characterized major molecular characteristics of R236C in two cellular platforms that served as an urothelial and a cholangiocyte-specific cell model. Both models were derived from primary cells of a patient encoding a novel mutation of ADPLD. The two cell models showed the expected individual characteristics of cells stemming from different organs. Aberrant functions caused by R236C were shared by the cells of the two models suggesting that the mutation, regardless of the cell type, has a dominant impact on proliferation, autophagy, apoptosis, cAMP synthesis, calcium homeostasis, and secretion. In summary, to our best knowledge, this is the first report on a novel channelopathy-associated mutant characterized side-to-side in two cell types having the same patient-specific genetic background.

The gold standard to study disease-causing mutations is commonly represented by animal models and patient-derived primary cells. However, with the exception of zebrafish [15] and blood-derived cells [16, 17] such models are not available for SEC61A1 channelopathies. For ADPKD, several patient-derived cell models have been established including short-time proliferating cells derived from kidney, immortalized cells, and iPSC-derived cell models [30]. Studies of SEC61A1 mutations are frequently performed in commercial cell lines (e.g. Hela, HEK293T) that are subjected to transient knockdown of wildtype *SEC61* followed by transgenic expression of the mutant [15–17]. Such expression systems lack the genuine genetic background of the patient and other organ- and cell-specific constraints. While a *PRKCSH* knockout in iPSCs was used to mimic ADPLD [31], our work describes for the first time the consequences of a PLD mutation in constitutively proliferating cells harboring the patient-specific genetic background. R236C-expressing cells showed stable cell growth which allowed various analyses difficult to obtain in short-time dividing primary tissue cell cultures. However, the proliferation of cells encoding R236C was reduced suggesting that the mutation affects check points of the regular cell cycle.

Two date, only six mutations of *SEC61A1* are known from patients, including 5 missense mutations. The missense mutation R236C, located in the third cytoplasmic loop [6], has a position that is quite distant from the other missense mutations [7]. For the other missense mutations of *SEC61A1*, major parameters of apoptosis, calcium homeostasis and protein translocation have been previously characterized. Autophagy was studied here for the first time using a *SEC61A1* mutation. Apoptosis was found to be upregulated in mutants V85D and Q32R, while translocation of proteins was reduced [16, 17, 32]. In addition, an altered calcium leakiness was observed in R236C-expressing cells in both cellular platforms. Such leakiness was also observed for V67G, T185A, V85D, and Q92R corroborating that various missense mutations may impact the passive transfer of calcium along the concentration gradient from the ER to the cytoplasm. It is thus tempting to speculate that basic cellular mechanisms disturbed in mutants of *SEC61A1* may play a role in the course of channelopathies which however cause various disease and affect different tissues, e.g. pneumonia, tumor, diabetes, common variable immunodeficiency (CVID), and as suggested here ADPLD [6, 10].

It is well established that cAMP has a key role in the regulation of fluid homeostasis and proliferation of epithelial cells [33]. In ADPKD patients and animal models of PKD/PLD increased levels of cAMP were observed [34–36]. In contrast, our findings indicate that R236C-expressing cells have a reduced intracellular level of cAMP suggesting that not all mutations causing ADPLD share a phenotype commonly observed in PKD. Use of somatostatin analogs that reduce cAMP levels has been proposed as standard-of-care drug treatment for PLD and clinical benefit was observed in a portion of PLD patients [37]. While our observation may not be representative for other ADPLD mutations, a detailed understanding of the cAMP homeostasis that modulate PLD seems to be important to identify appropriate therapeutic targets

Autophagy and apoptosis are critical for the functional integrity of organs and cells under normal and pathological conditions and, although distinctly regulated, can be closely related [38]. One central finding of our study is that autophagy is reduced, while apoptosis, upon oxidative stress, is increased in R236C-expressing cells. A reciprocal relationship between apoptosis and autophagy was noted before in murine models of PKD, where suppressed autophagy is accompanied by increased apoptosis associated to proliferation and cyst formation [26, 39]. A reduced autophagy was mostly observed in PKD models and a few patients [39–41]. However, the opposite was observed in other models and some ADPKD patients suggesting that methodology of autophagy determination and peculiarities of the model may impact results [42–44].

Our results of a reduced autophagy in R236C-expressing cells were confirmed by two independent methodologies in both cell lines using western blot and flow cytometry analysis suggesting that SEC61A1 mutation R236C might differ from mutations that cause ADPLD. With regard to apoptosis, an increase of apoptotic markers was observed in PLD liver sections [45–47]. R236C-expressing cells showed normal apoptosis under basal conditions as observed in PKD [48], while at a condition of oxidative stress, cells were highly prone to apoptosis and necrosis. Enhanced apoptosis was also observed in the kidney ADPKD [45, 49–51] suggesting that oxidative stress, commonly associated with inflammatory responses, could have a common role in tissue damage of PKD and PLD [52].

The establishment of liver cysts is thought to be dependent on multiple events, including a disease-causing mutation (primary), a secondary (unknown) somatic mutation, and progressive functional abnormalities of excessive fluid secretion and enhanced cyst proliferation (tertiary) [3]. Although the understanding of the molecular mechanisms operative in PKD and PLD has significantly improved over the last decades, the exact mutual dependences of the mechanisms for development of disease remain obscure. Our data provide first evidence that SEC61A1 mutation R236C shows several perturbations of the mechanisms proposed as being hallmarks of polycystic disease. Of note, individual transgenes and cell sources used for establishment of cell models may impact results [30]. In consideration of such effects, our observations were recorded in two different cell types established by two different genetic methodology. However, with regard to the urine-related cell source that was used for both models, it is likely that additional disease-modifying, secondary somatic mutations were not implemented, as such mutations are proposed to be only present in some regions of the liver. Apart from the characterized first mutation, the secondary and tertiary events orchestrating the development of polycystic disease specifically in the liver but not in other organs are still far from being understood. In summary, we have established patient-specific cell platforms by two different methodology to understand the molecular phenotype of a novel mutation observed in ADPLD. Such methodology may allow to decipher the pathophysiologic mechanisms of mutations identified in the genetically diverse group of PLD patients and in other channelopathies to guide molecularly targeted therapy in the future.

## Acknowledgement

Part of the work was funded by the Ministerium für Kultur und Wissenschaft des Landes Nordrhein-Westfalen (PtJ-Az. 2305st004). The authors declare no competing financial interests.

